# Validity and reliability of a new whole room indirect calorimeter to assess metabolic response to small-calorie loads

**DOI:** 10.1101/2023.09.21.558672

**Authors:** Mary Elizabeth Baugh, Monica L. Ahrens, Zach Hutelin, Charlie Stylianos, Erica Wohlers-Kariesch, Mary E. Oster, Jon Dotson, Jon Moon, Alexandra L. Hanlon, Alexandra G. DiFeliceantonio

**Affiliations:** Fralin Biomedical Research Institute at VTC, Roanoke, VA; Center for Health Behaviors Research at Fralin Biomedical Research Institute at VTC, Roanoke, VA; Center for Biostatistics and Health Data Science, Department of Statistics, Blacksburg, VA; Translational Biology, Medicine, and Health, Fralin Biomedical Research Institute at VTC, Roanoke, VA; Department of Human Nutrition, Foods, and Exercise, Virginia Tech, Blacksburg, VA; MEI Research, Ltd, Edina, MN

**Keywords:** Metabolic chamber, macronutrient oxidation, validation, diet induced thermogenesis, carbohydrate

## Abstract

**Objective:** To provide an overview of our whole room indirect calorimeter (WRIC), demonstrate validity and reliability of our WRIC, and explore a novel application of Bayesian hierarchical modeling to assess responses to small carbohydrate loads.

**Methods:** Seven gas infusion studies were performed using a gas blender and profiles designed to mimic resting and postprandial metabolic events to assess WRIC validity. In a crossover design, 16 participants underwent fasting and postprandial measurements, during which they consumed a 75-kcal drink containing sucrose, dextrose, or fructose. Linear mixed effects models were used to compare resting and postprandial metabolic rate (MR) and CO (CO). Bayesian Hierarchical Modeling was also used to model postprandial CO trajectories for each participant and condition.

**Results:** Mean total error in infusions were 1.27 ± 1.16% and 0.42 ± 1.21% for VO_2_ and VCO_2_ respectively, indicating a high level of validity. Mean resting MR was similar across conditions (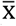 =1.05 ± 0.03 kcal/min, *p*=0.82, ICC: 0.91). While MR increased similarly among all conditions (∼13%, *p*=0.29), postprandial CO parameters were significantly lower for dextrose compared with sucrose or fructose.

**Conclusions:** Our WRIC validation and novel application of statistical methods presented here provide important foundations for new research directions using WRIC.

## Introduction

Indirect calorimetry is a valuable tool to assess metabolic response to various physiological stimuli or interventions. Due to technological advancements in recent years in instrument sensitivity and data processing techniques, interest in measuring small, dynamic changes in energy expenditure and macronutrient oxidation following short exercise bouts or small dietary intake loads is increasing.

Historically, whole-room indirect calorimeters (WRICs) have required long periods of time (i.e., several hours) to quantify O_2_ consumption and CO_2_ production due to large room volumes and lower sensitivity of measurement equipment (1); however, advancements in WRIC hardware over recent decades have improved response time and resolution. While metabolic carts have traditionally been used to assess metabolic changes over short or dynamic measurement periods, some inherent limitations constrain their use, including 1. participant discomfort from facemask or canopy placement, 2. gas analyzer drift during longer measurements (2), and 3. between-instrument reliability and reproducibility (3). Small-volume WRICs have the potential to overcome these limitations while also providing advantages over large-volume WRICs in terms of reduced room volume and rate of room air turnover.

The thermic effect of food (TEF) is estimated to comprise approximately 10% of total daily energy expenditure (4); however, it can vary widely based on the macronutrient composition of a meal (5). Specifically among carbohydrates, the greater increases in TEF and carbohydrate oxidation (CO) after ingestion of large amounts (i.e., 300 kcal) of fructose and sucrose, which contains fructose, compared with dextrose and dextrose polymers have been well characterized (5–9). However, whether TEF or CO responses differ among smaller carbohydrate loads has not been thoroughly assessed. Given that sugar sweetened beverages make up the majority of added sugars in the average American diet and contain 140-150 kcals/serving on average (10), testing the metabolic response to carbohydrate loads <300 kcals is an important research gap.

Common statistical methods to assess TEF include calculations of area under the curve (AUC) followed by comparisons of the AUCs across groups (11, 12). However, these methods are limited in that they condense serial measurements to a single summary parameter before statistical comparisons are performed, which introduces greater variance and decreases power. In addition, physiologically meaningful information can be lost. For example, two different curves can have the same total area under the curve (AUC) but different trajectories over time; only comparing AUCs would fail to capture true differences in metabolic responses. One alternative method is Bayesian Hierarchical Modeling, which accounts for differing temporal trajectories across observations by utilizing multiple parameters to estimate individual smoothed curves for each measure of interest across time and can also minimize the variance for summary parameters.

Therefore, our overall objectives were to: 1. provide a methodological overview of our small-volume WRIC system and evidence of instrument validity and reliability; 2. demonstrate reliability in measuring physiological variables in human studies; and 3. explore the temporal resolution of metabolic responses elicited by small carbohydrate loads (i.e., 75 kcals) using a novel application of Bayesian Hierarchical Modeling.

## Materials and Methods

### Whole Room Indirect Calorimeter Description

The small-volume WRIC (MEI Research, Ltd) at the Fralin Biomedical Research Institute at Virginia Tech Carilion, built in 2019, is a 1.2 x 2.1 x 2.3 m WRIC designed for both resting and exercise measurements. WRIC volume for the resting configuration was 4730 L, determined by washout tests (**Figure 1**). Inflow air to the WRIC is provided by a medical air system separate from the building supply, which minimizes the influence of diurnal fluctuation in gas concentration of atmospheric air. The chamber has the capability of being operated in either “push” or “push-pull” modes. When operating in “push-pull” mode, an air blower provides a vacuum on the outflow MFC, which regulates pressure inside the chamber. This allows for control of both inflow and outflow rates so that the room can be operated with a lower ventilation rate and at a minimal pressure difference. The present study was operated in “push” mode, in which the medical air system “pushes” inflow air, creating a positive pressure inside the WRIC. Proportional-integral-derivative (PID) control of air inflow rate adjusts to control CO_2_ concentration inside the WRIC to a constant and optimal range of measurement for the analyzer. Inflow and outflow O_2_ are continuously measured with a dual channel paramagnetic sensor (Siemens Oxymat6), and inflow and outflow CO_2_ are continuously measured with an infrared sensor (Siemens Ultramat6). The O_2_ analyzer has a constantly flowing reference from a gas tank (∼21% O_2_, balance N_2_). The CO_2_ analyzer has a sealed reference cell filled with N_2_. Perma Pure dryers (Perma Pure, LLC, PD-50T-48MSS) remove water vapor from sample gases prior to entering outflow analyzers, and humidity sensors (Viasala, HMP60C12A0A3B0) verify adequately dried samples. In addition, temperature (Vaisala, HMP60C12A0A3B0), humidity (Vaisala, HMP60C12A0A3B0), and pressure (Omega, PX653-2.5BD5V) inside the WRIC are monitored by specific sensors.

**Figure 1:**
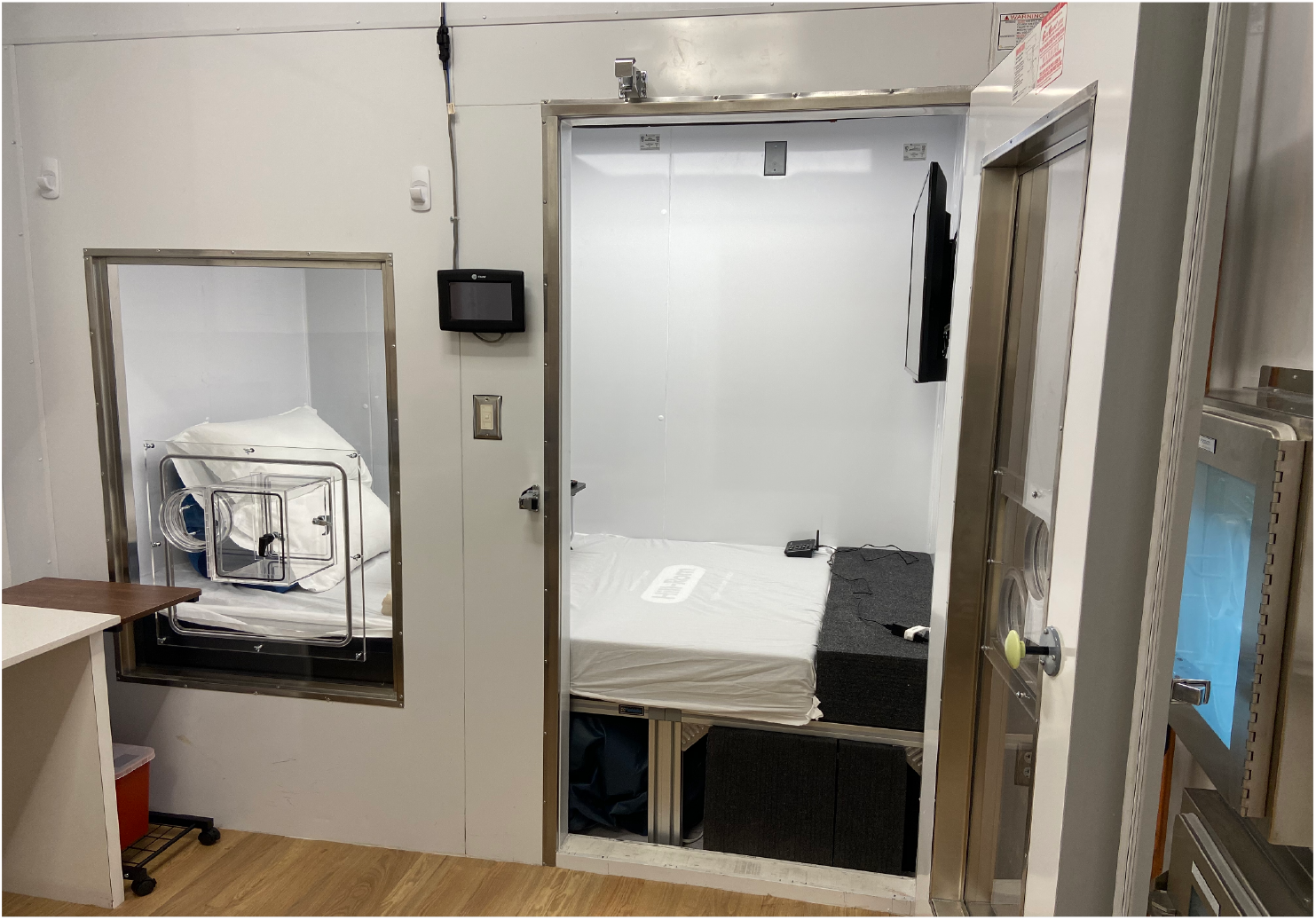
The small-volume “flex” metabolic chamber at the Fralin Biomedical Research Institute at Virginia Tech Carilion in Roanoke, Virginia, set up for a resting measurement.

### Routine Calibrations

After each O_2_ analyzer reference gas tank change, a hardware calibration is performed on the O_2_ analyzer, in which the absolute measurement range of the analyzer is established.

Following the hardware analyzer calibration, a blender calibration is performed to ensure linearity of the O_2_ analyzer. During the blender calibration process, a gas blender is used to gflow O_2_, CO_2_, and N_2_ through the full range of the analyzers. Linearity of the O_2_ analyzer readings is then verified (r^2^>0.9999), and differences between known and measured gas concentrations are then calculated to be applied as corrections to VO_2_ and VCO_2_ data collected during experiments. While the CO_2_ analyzer is not inherently linear, linearity of this analyzer is also verified (r^2^>0.999).

### Bi-Annual Maintenance

All high flow and blender mass flow controllers (MFCs; Alicat Scientific) are calibrated bi-annually by MEI, Ltd. using positive displacement primary piston provers specific to each MFC (ML-800 and ML-1020, Mesa Labs). In the calibration process, each MFC is first tested at 10 points across the full flow range, beginning at 10%; a second test across the full range is then run at 10% intervals beginning at 5%. Each MFC must measure within 0.5% of the prover to pass calibration. All blender MFCs are calibrated using their respective gasses (i.e., N_2_, O_2_, and CO_2_).

### Validation Studies

#### Infusion Validation Studies

Overall system is validated regularly using blender infusions of dry N_2_ and CO_2_ in two protocol profiles designed to test detection of minimal changes in gas concentrations (i.e., test limits of detection within a physiologically relevant range) as well as mimic anticipated resting and postprandial measurements for the present human study design.

#### Metabolic Chamber Data Post-Processing

Measured inflow and outflow O_2_ and CO_2_ concentrations for all infusion and human studies were adjusted by linear interpolation using corrections established during blender calibrations (see Supplemental Methods). Calculated physiological variables of interest (e.g., metabolic rate (MR), respiratory exchange ratio (RER), etc.) and variables to ensure measurement validity (e.g., chamber pressure, temperature, etc.) were recorded using CalRQ (MEI Research, Ltd), a customized software developed in LabVIEW. VO_2_ and VCO_2_ were calculated using standard equations incorporating WRIC volume, fractional concentrations of O_2_ and CO_2_ of inflow air and WRIC air, and inflow and outflow rates. After collection, an 8-minute centered derivative term was applied to data during post-processing to determine VO_2_ and VCO_2_.

### Human Studies

#### Human Study Ethics Approval

The human study protocol was approved by the Virginia Tech Institutional Review Board (#21-052). All participants provided verbal and written informed consent prior to participation in the study.

#### Participants and Experimental Design

Sixteen males and females without metabolic disease completed the study from July 2021-October 2022. Participants were not adhering to specific dietary patterns (e.g., intermittent fasting, ketogenic/low-carb diets) prior to enrolling. Participants reported weight stability (i.e., weight change ≤5 lbs) for the previous 3 months and reported not taking medications known to influence study measures, including antiglycemic agents, thyroid medications, and sleep medications. They also reported not using tobacco or nicotine products.

After a consent visit, which included anthropometric measurements of height, weight, and waist and hip circumference, participants completed 3 separate WRIC sessions in a randomized crossover design. On the day of each metabolic chamber session, participants reported to the laboratory between 6 and 9 am after fasting for ≥8 h. Participants were also instructed to abstain from physical activity for 12 h prior to each session and to consume a specified meal meeting 35% of their estimated energy requirements between 6-10pm the evening before each session. After a 45-min resting measurement inside the chamber, participants were instructed to consume a 355 ml beverage containing 75 kcal of either sucrose, dextrose, or fructose. Beverages were made by mixing the carbohydrate with a solution of deionized water, flavoring (Bell Labs), and food coloring (McCormick & Co). Participants were blinded to drink composition and were instructed to consume the beverage within 5 minutes and immediately resume resting for the duration of the measurement. Post-beverage indirect calorimetry measurements were collected continuously for approximately 80 minutes (range: 55-111 minutes). Some measurements lasted <80 minutes (n=10) due to compliance and technical issues. Participants were allowed to watch TV, read, or listen to music or podcasts while inside the WRIC.

#### Control of Dietary Intake

During a consent visit, estimated energy needs were calculated for each participant using the Mifflin-St. Jeor equation multiplied by an activity factor based on self-reported exercise and physical activity patterns (Mifflin 1990). Participants were then instructed to consume a specified meal (50% of energy from carbohydrate, 30% from fat, and 20% from protein) meeting 35% of their estimated energy requirements between 6-10pm the evening before each session. Foods included in the meal were individualized to participant preferences but contained the same macronutrient composition.

When participants arrived at the laboratory for each experimental session, a 24-hour dietary recall for the previous day was collected using a multiple pass method (13, 14); this recall included the specified meal planned during the consent visit. Dietary recalls were analyzed using Nutrition Data Systems for Research (version 2020).

#### Statistical Analysis

Energy expenditure was calculated using the modified Weir equation (15), and substrate oxidation was calculated using fat and CO equations (16) based on published respiratory quotient (RQ) tables (17). Data for the 4 minutes immediately before and after participants drank the test beverage were excluded to account for participant movement during beverage consumption that would have been captured by an 8-minute centered derivative applied during post-processing. Thus, baseline measures were calculated as the average of minutes -20 to -4 preceding beverage consumption. Mean fasting time, dietary intake, and RMR were compared across the three conditions using linear mixed effects models (LMM). Additionally, intra-class correlation coefficients (ICCs) were estimated from the LMMs. We interpreted an ICC value >0.75 as indicating very good reliability. Due to the lack of detectable differences across conditions in fasting time, dietary intake variables, or RMR, we proceeded with our other analyses without accounting for these variables.

Two statistical analysis methods, the commonly-used LMM method and a Bayesian Hierarchical Model approach, were used. The LMM method included testing for differences in time until peak CO, peak CO, and area under the curve (AUC) for change in CO across the three conditions. AUC was calculated using the trapezoidal rule; for measurements lasting <80 minutes (n= 10), the last measurement value was carried forward through 80 minutes.

Our proposed method used a Bayesian Hierarchical Model approach to model the CO trajectories for each subject and condition. With estimates from this model, tests to compare AUC, peak CO, and time at peak CO were conducted. Establishing this model has two main benefits in that it (1) uses all data points in the model, allowing more statistical power to detect differences across groups compared with the traditionally used LMM method; and (2) smooths each participant’s data, allowing for a well-defined maximum value and return to baseline. The model assumes that the average difference in CO from resting across time takes the functional form of a natural cubic spline,

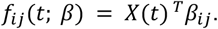

Where *X*(t) is the basis matrix for the natural cubic spline with knots at time points 20 and 48. These knots were chosen as two derivatives after the minimum time point and two derivatives before the maximum observed time point. To allow for measurement error, we assumed that data follows a normal distribution with mean of the cubic spline function, *f*(*t*; *β*_*ij*_), and variance of *σ*_*e*_:

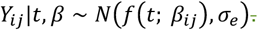

Where *Y*_*ij*_ is the outcome of interest – percent change from resting CO. In this model, *β*_*ij*_ corresponds to the coefficient estimate for subject *i*’s *j*^th^ condition,

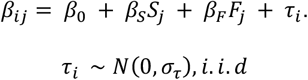

Where *β*_0_ is the vector of coefficients for the dextrose condition, *β*_*s*_ is the change of the coefficients for the sucrose condition relative to the dextrose condition, and *β*_*F*_ is the change of the coefficients for the fructose condition relative to the dextrose condition. *S*_*j*_ is an indicator function for condition *j* being the sucrose and *F*_*j*_ is an indicator for condition *j* being the fructose, *j =* 1, 2, 3. Finally, τ_*i*_ is the individual intercept, which accounts for the within-subject design of the study. All priors were chosen to be non-informative to let the data dictate the estimates for all unknown coefficients. The statistics of interest were estimated using the estimated values from fitting the Bayesian model using:

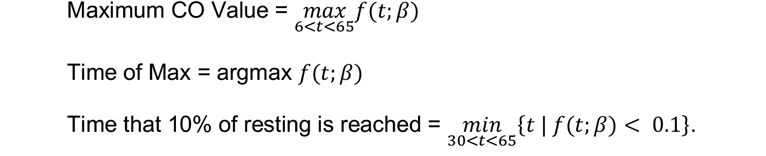

These statistics were estimated using the mean from the approximate posterior distribution. To compare these statistics across conditions, 95% credible intervals of the posterior distribution for the differences are reported. The 95% Bayesian credible interval is analogous to the frequentist 95% confidence interval, in that a 0 contained within the interval indicates no statistically significant difference between conditions.

The benefit of this Bayesian approach is that it allows a flexible model to be fit using only 16 participants. Each participant has their own mean structure estimate for each condition, which allows an estimate of the three statistics of interest on both individual and group levels. Additionally, the method results in estimates of each parameter’s distribution, allowing for easy statistical comparison between groups. The R package, Nimble, was used to fit this model (18). Nimble is a Markov chain Monte Carlo sampler that uses Gibbs sampling (18) to get approximate posterior distributions of all parameters in the defined model. Data are reported as means and standard deviations, unless otherwise specified; statistical significance for the traditional LMM models were set at *p*<0.05.

## Results

### Validation Studies

Over all infusions spanning the length of human data collection in this study (n=7), mean total error in VO_2_ and VCO_2_ were 1.27 ± 1.16% and 0.42 ± 1.21%, respectively. To assess reproducibility of gas recoveries across specific gas analyzer ranges that mimic the anticipated human study measurements, infusions were also analyzed by section. A sample infusion tracing expected and observed gas recoveries is shown in **Figure 2**. Results for each section of one profile type (n=3) are shown in **Table 1**. Error for VO_2_ and VCO_2_ ranged -0.97 ± 1.13% to -1.22 ± 1.27% and 0.42 ± 0.58% to 1.04 ± 1.40%, respectively, across sections. Over all infusions (n=7), gas recovery rates were 98.8 ± 1.2% for VO_2_ and 100.5 ± 1.2% for VCO_2_.

**Figure 2.**
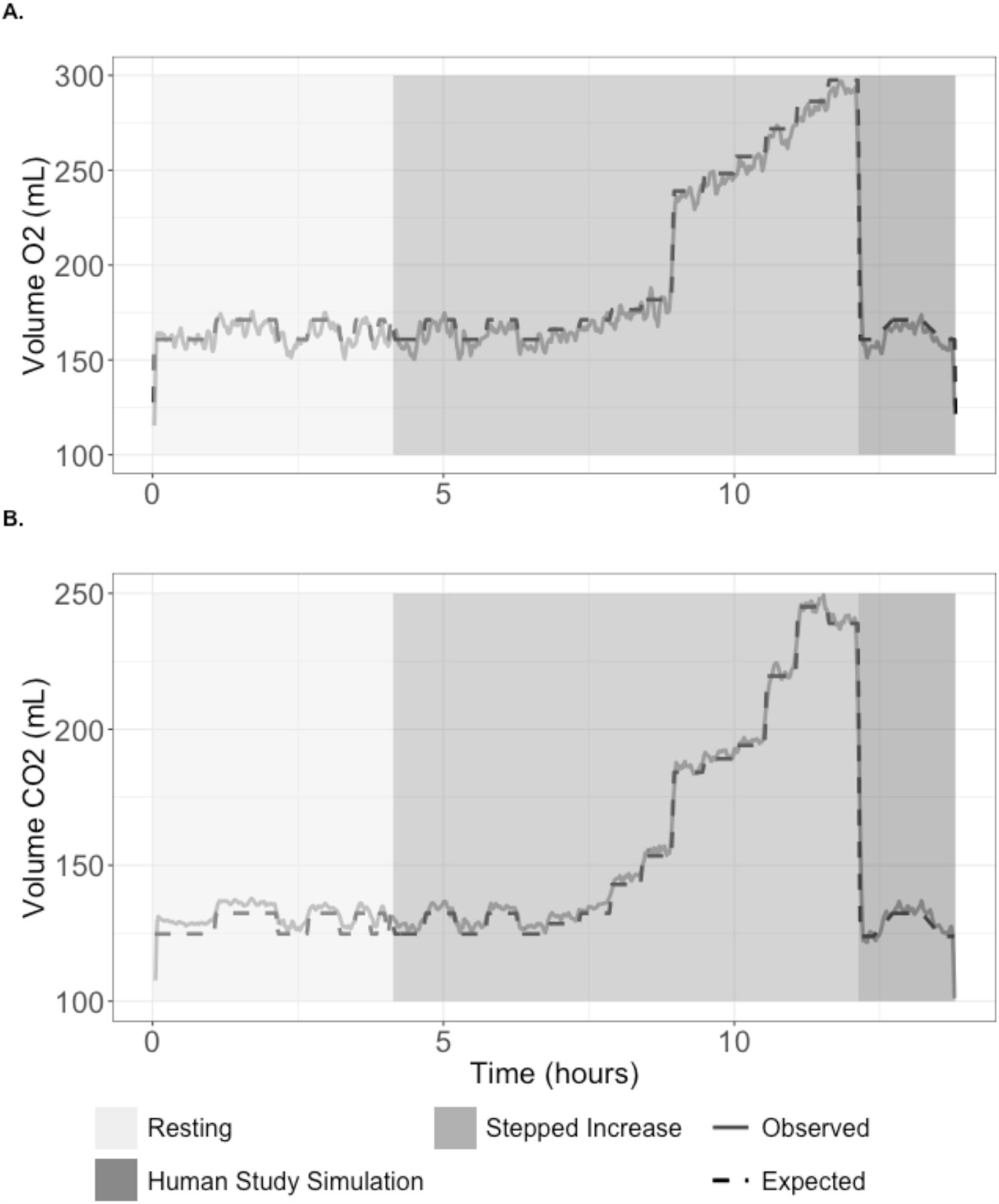
A sample infusion depicting (dashed) expected and (solid) observed (A) VO_2_ and (B) VCO_2_ values. Resting, stepped increase, and human study simulation event sections are denoted by differing background shading.

**Table 1.** Reproducibility of O_2_ and CO_2_ recovery across specified event sections during infusion validation studies (n=3)

| Event | $\Delta VO_2$<br>(expected – observed)<br>(mL/min) | Error<br>$VO_2$ (%) | MDC<sup>a</sup> $VO_2$<br>(mL/min) | $\Delta VCO_2$<br>(expected – observed)<br>(mL/min) | Error<br>$VCO_2$ (%) | MDC<sup>a</sup> $VCO_2$<br>(mL/min) |
| --- | --- | --- | --- | --- | --- | --- |
| Resting | 1.87 ± 2.5 | -1.13 ± 1.5 | 1.03 ± 0.09 | -1.46 ± 1.8 | 1.14 ± 1.4 | 0.73 ± 0.05 |
| Stepped Increase | 2.57 ± 2.66 | -1.24 ± 1.28 | 0.65 ± 0.26 | -0.8 ± 0.99 | 0.49 ± 0.6 | 0.43 ± 0.23 |
| Human Study Simulation | 1.62 ± 1.9 | -0.99 ± 1.15 | 2.31 ± 0.43 | -1.06 ± 0.52 | 0.84 ± 0.4 | 1.77 ± 0.31 |
| Total | 2.24 ± 2.49 | -1.18 ± 1.31 | 0.56 ± 0.14 | -1.03 ± 1.15 | 0.69 ± 0.77 | 0.4 ± 0.11 |

<sup>a</sup>MDC, minimal detectable change was calculated as standard error of the mean x 1.96 x $\sqrt{2}$ , where 1.96 is the critical value and $\sqrt{2}$ is the correction factor for measurement in duplicate.
Data are expressed as mean ± standard deviation.

### Control of Potential Biological Confounders

Participant characteristics are shown in **Table 2**. Fourteen females and 2 males aged 29 ± 6 years with a body mass index 24.3 ± 4.4 kg/m^2^ completed the study. Mean fasting time prior to WRIC session was not different among groups (*p* = 0.19). On average, participants fasted approximately 11.3 ± 0.2 hours before measurements. Self-reported dietary intake was not different across carbohydrate conditions for total energy, fat, carbohydrate, or protein intake (**Supplemental Table 1**). Participants reported consuming ∼33-36% of kcals from fat, 45-49% of kcals from carbohydrate, and 17-20% of kcals from protein during the 24 hours prior to each WRIC session, which reflects the macronutrient composition of the prescribed evening meal.

**Table 2.** Participant Characteristics (n=16)

|  | Mean (SD) or n (%) |
| --- | --- |
| **Age (years)** | 29 (6) |
| **Sex** |  |
| Female | 14 (87.5%) |
| Male | 2 (12.5%) |
| **Race** |  |
| White | 11 (68.8%) |
| White and other | 3 (18.8%) |
| Black or African American | 2 (12.5%) |
| **Ethnicity** |  |
| Hispanic, Latino, or Spanish Origin | 1 (6.3%) |
| Non-Hispanic, Latino, or Spanish Origin | 15 (93.7%) |
| **Body weight (kg)** | 69.2 (14.4) |
| **Habitual Dietary Pattern** |  |
| Vegan | 1 (6.3%) |
| Vegetarian | 2 (12.5%) |
| **BMI (kg/m<sup>2</sup>)** | 24.3 (4.4) |
| **Waist-to-hip ratio** | 0.77 (0.07) |
| **Estimated RMR (Mifflin-St. Jeor) (kcal/d)** | 1449 (181) |
| **Measured RMR (kcal/d)** | 1483 (216) |

BMI, body mass index; RMR, resting metabolic rate

### Metabolic Rate

Mean RMR, as estimated by LMM, was similar across all conditions (*p*=0.82; **Figure 3a**), and the ICC across all three conditions was 0.91, indicating good test-retest reliability within participants. Mean RMR across conditions was 1.05 ± 0.03 kcal/min. Change in MR in response to conditions is depicted as the elevation in MR above resting (**Figure 4**); MR increased similarly after all carbohydrate loads, and calculated AUCs for change in MR did not differ among carbohydrate types (*p*=0.29; **Figure 4a**). In addition, among all conditions MR increased by 13% from baseline (*p*<0.001).

**Figure 3.**
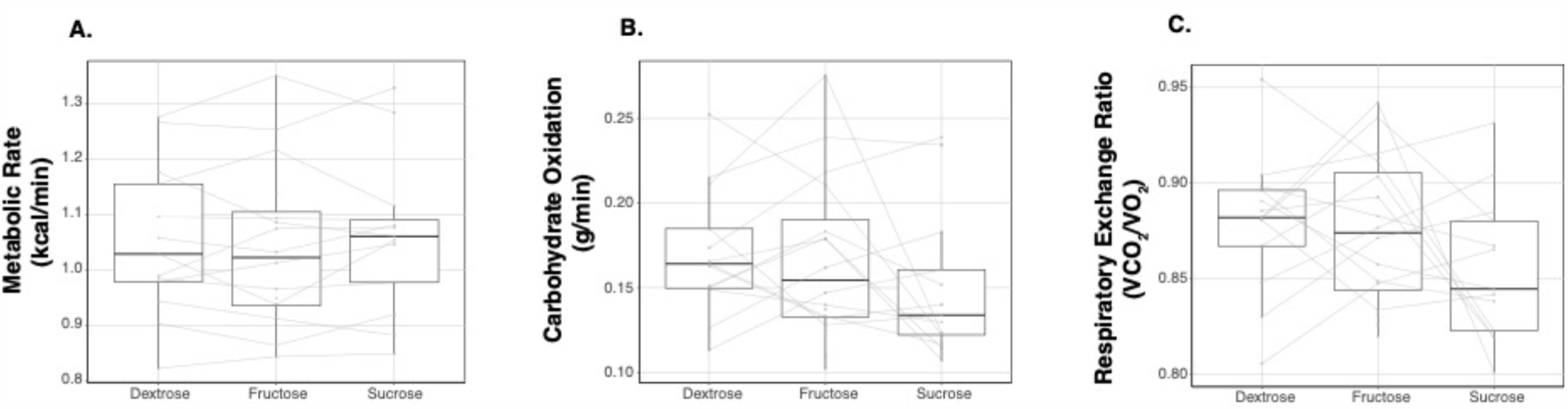
Mean resting (A) metabolic rate, (B) carbohydrate oxidation, and (C) respiratory exchange ratio across dextrose (n=13), fructose (n=16), and sucrose (n=13) conditions.

**Figure 4.**
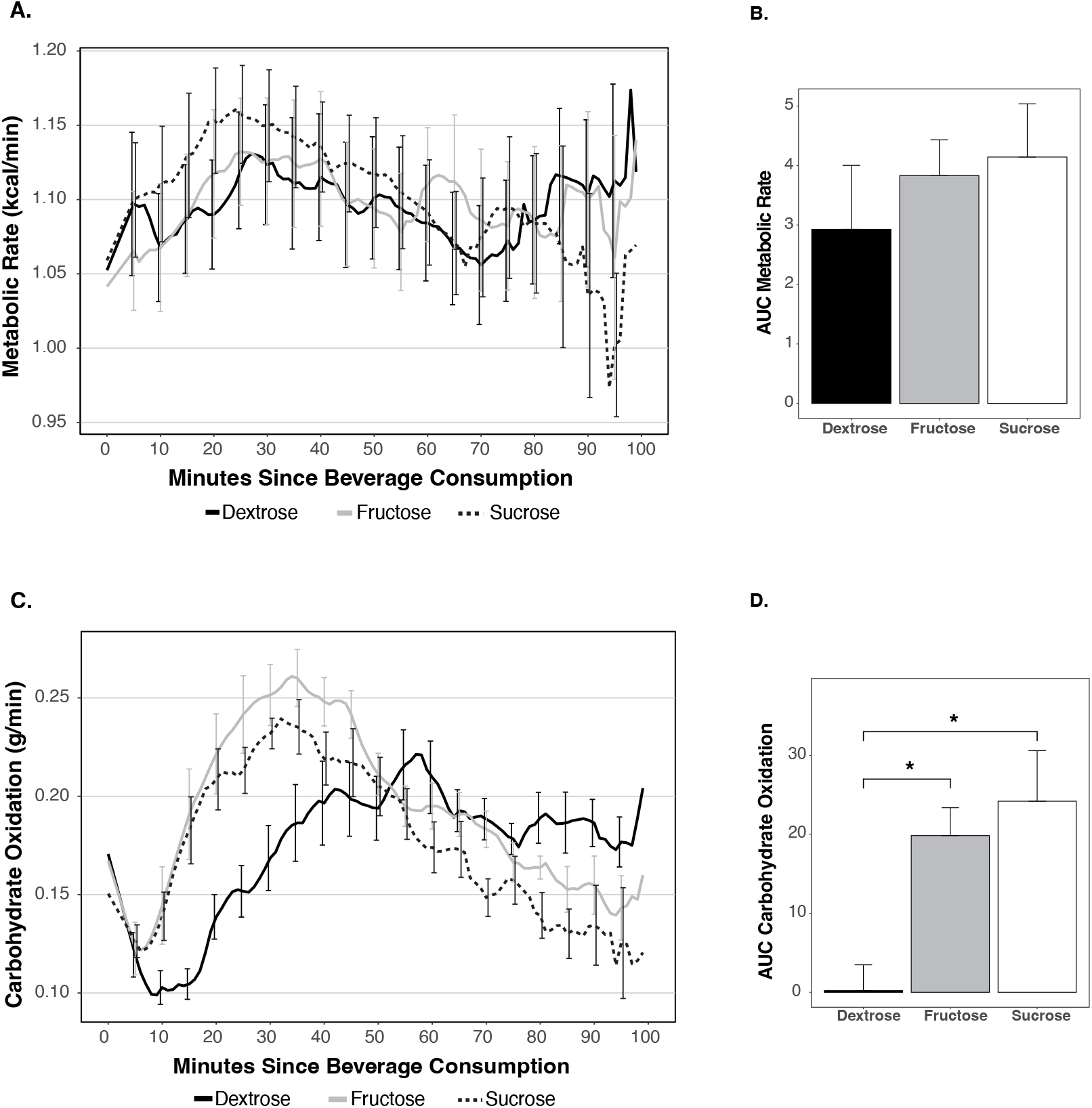
Postprandial change from resting (A) metabolic rate and (C) carbohydrate oxidation and areas under the curve for percent change from resting (B) metabolic rate and (D) carbohydrate oxidation in response to consumption of 75-kcal beverages containing dextrose, fructose, or sucrose. Data are expressed as mean ± standard error of the mean.

### Substrate Oxidation

Mean resting RER and CO were not different across conditions (*p*=0.15 and *p*=0.26, respectively; **Figures 3b and 3c**), but the intraclass correlation coefficients were 0.004 and 0.324 for RER and CO measures across conditions, respectively, suggesting weak to moderate reliability across test days. Peak CO was lower after dextrose (0.24 ± 0.07 g/min) compared with fructose (0.29 ± 0.06 g/min; p < 0.01) and sucrose (0.27 ± 0.04 g/min; p = 0.06) consumption. Time to reach peak CO was longer after consumption of the dextrose (50.8 ± 12.1 minutes) compared with fructose (35.4 ± 10.8 minutes; p < 0.01) or sucrose (33.5 ± 10.4 minutes; p < 0.01) beverages. There were statistically significant differences in calculated AUCs for change in CO from baseline among conditions. Change in CO AUC was significantly smaller for dextrose (-0.27 ± 1.75 g/min/min) compared with fructose (2.42 ± 1.74 g/min/min; p < 0.01) and sucrose (2.72 ± 2.26 g/min/min; p < 0.01).

### Bayesian Model

The Bayesian model resulted in similar mean estimates of CO peak change, time to reach peak, and AUC, but the variance in the estimates were lower than that from the standard methodology (**Table 3**). Time to reach peak CO was longer after consuming dextrose (57 min) compared with fructose (33 min, 95% Bayesian Credible Interval (BCI) [22, 27]) or sucrose (32 min, 95% BCI [24, 28]). Peak percent change in CO from resting mean was lower after dextrose (26%) compared with fructose (57%, 95% BCI [27, 35]) and sucrose (69%, 95% BCI [39, 47]) consumption. There were statistically significant differences in calculated AUCs for percent change in CO from baseline among conditions. Change in CO AUC was significantly lower for dextrose (4.4) compared with fructose (22.4, 95% BCI [16.1, 19.8]) and sucrose (30.5%, 95% BCI [24.2, 28.0]).

**Table 3.**
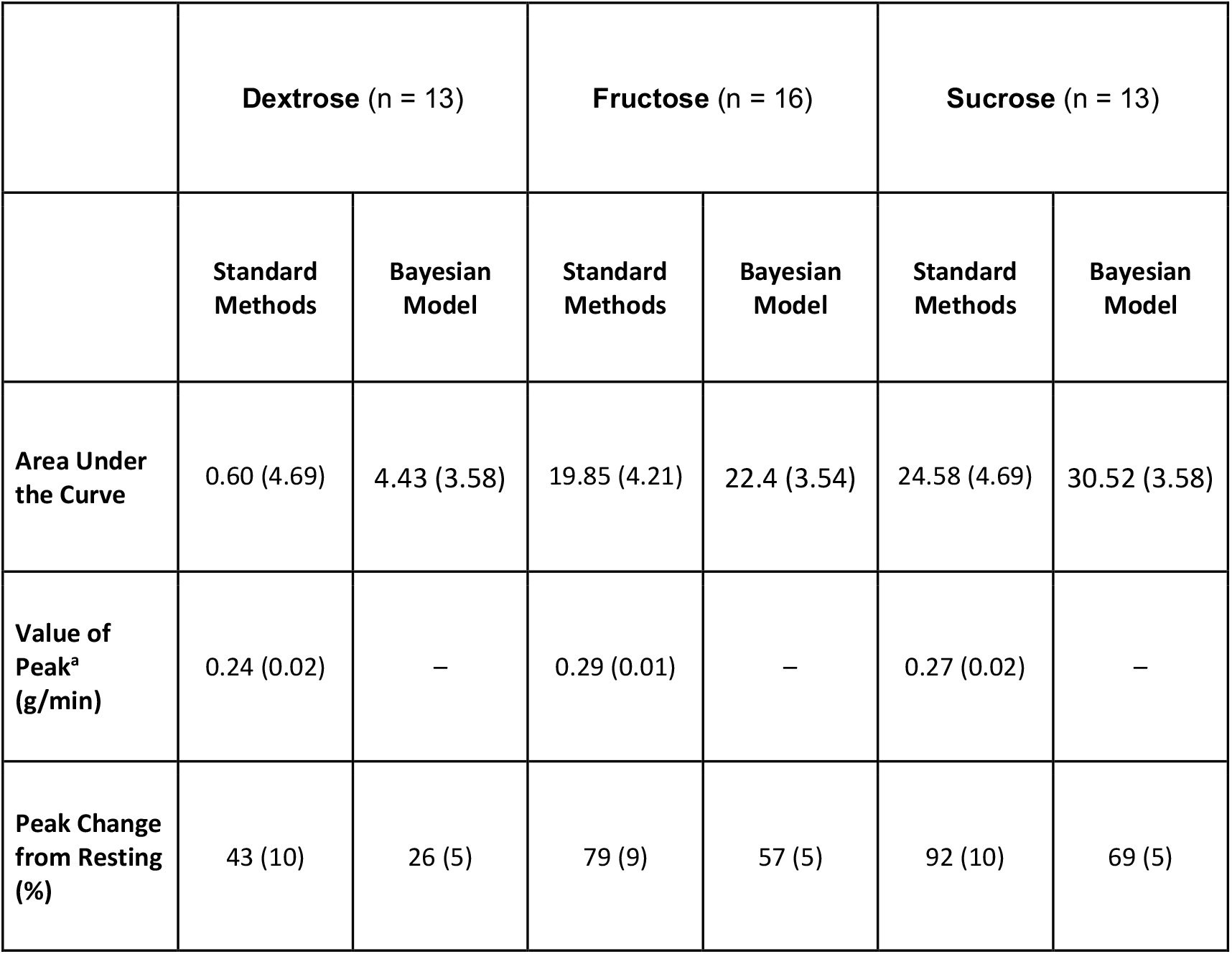
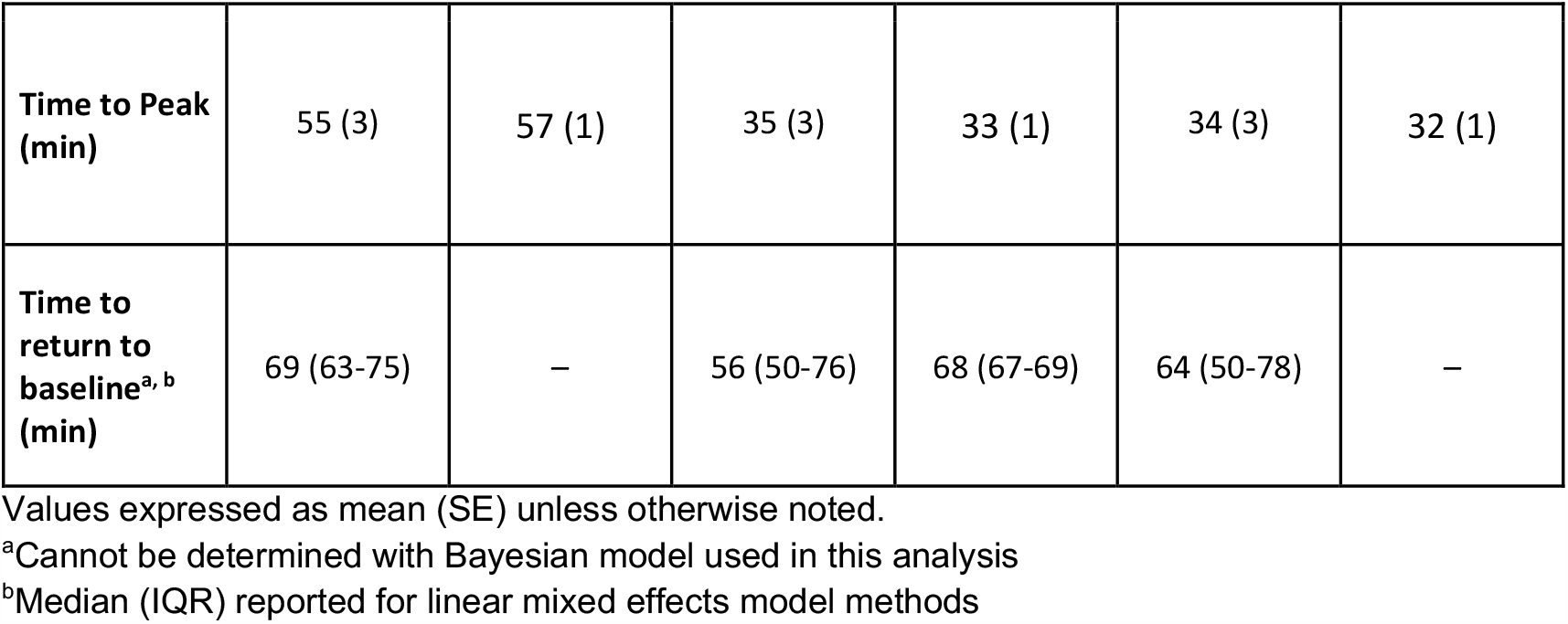
Comparison postprandial carbohydrate oxidation parameters in response to dextrose-, fructose-, and sucrose-containing beverage consumption as assessed by standard methods and Bayesian hierarchical modeling statistical methods.

|  | Dextrose (n = 13) |  | Fructose (n = 16) |  | Sucrose (n = 13) |  |
| --- | --- | --- | --- | --- | --- | --- |
|  | Standard Methods | Bayesian Model | Standard Methods | Bayesian Model | Standard Methods | Bayesian Model |
| Area Under the Curve | 0.60 (4.69) | 4.43 (3.58) | 19.85 (4.21) | 22.4 (3.54) | 24.58 (4.69) | 30.52 (3.58) |
| Value of Peak<sup>a</sup> (g/min) | 0.24 (0.02) | – | 0.29 (0.01) | – | 0.27 (0.02) | – |
| Peak Change from Resting (%) | 43 (10) | 26 (5) | 79 (9) | 57 (5) | 92 (10) | 69 (5) |
| **Time to Peak (min)** | 55 (3) | 57 (1) | 35 (3) | 33 (1) | 34 (3) | 32 (1) |
| **Time to return to baseline<sup>a, b</sup> (min)** | 69 (63-75) | – | 56 (50-76) | 68 (67-69) | 64 (50-78) | – |

Values expressed as mean (SE) unless otherwise noted.
<sup>a</sup>Cannot be determined with Bayesian model used in this analysis
<sup>b</sup>Median (IQR) reported for linear mixed effects model methods

## Discussion

A recent report outlines essential guidelines for reporting WRIC data (4), which include providing methods for standardizing data reported and evidence for validity of the WRIC system collecting the data. Following these guidelines in reporting and research publications will ultimately allow for comparisons across calorimeters and institutions regardless of equipment type and validation methods. Using these guidelines, we sought to provide evidence of the validity and reliability of a small-volume WRIC at the Fralin Biomedical Research Institute at Virginia Tech Carilion and to demonstrate accuracy in assessing small, dynamic changes in human respiratory VO_2_ and VCO_2_ in response to small calorie loads. Overall, results from our technical validation studies indicate a high degree of precision and accuracy in our system for detection of both resting and small, dynamic changes in gas concentrations. Results from our human studies confirm reliability in our measurements and provide evidence that differences in metabolic responses to small-calorie carbohydrate loads can be detected using WRIC.

### System Validation and Technical Reliability

Our results indicate high accuracy in recovery rates of VO_2_ and VCO_2_ and high reproducibility across multiple dry gas blender infusion studies. Though few studies have reported system validation data in publications, our overall recovery rates (98.9% for VO_2_ and 100.5% for VCO_2_) are comparable to rates reported by others (19, 20). To explore the validity of our system across a wider range of metabolic responses elicited by our human research study designs, we also assessed error rates for different “sections” of infusion studies. Error rates across each of these sections were <2% and minimal detectable changes ranged 0.42-2.31 mL/min, suggesting a high degree of measurement accuracy in VO_2_ and VCO_2_ response during both resting and short-duration, dynamic changes.

### Human Study Reliability

In our crossover design human study, mean RMR, resting RER, and resting CO were not different across conditions. In addition, mean dietary intakes for the 24 hours preceding measurements were not different across conditions. Taken together, these physiological findings reflect the results of technical validation studies indicating good reliability of our WRIC system. However, we also observed a high amount of intraindividual variability in measures of RER and CO across days. This is not unexpected, as RER is a ratio of VCO_2_ and VO_2_ and, thus, small divergences in gas concentration values will mathematically result in larger discrepancies in RER values. Furthermore, fasting RER values have been shown to primarily depend on the food quotient of dietary intake in previous days (21). Though we attempted to provide some control over dietary intake by standardizing macronutrient composition of the previous evening’s meal, our methods may not have been rigorous enough to control fasting RER and CO values. Others also have noted similar discrepancies in calculated RER and macronutrient oxidation values across repeated measures (4, 22, 23).

### Dynamic Metabolic Response Following a Small-Calorie Load

The thermic effect of food, defined here as the elevation in metabolic rate above resting, was not different among carbohydrate types. On average, MR increased 13% after consumption of a 75-kcal carbohydrate beverage. Previous studies assessing temporal response of MR to various carbohydrate loads have reported greater change in MR in response to fructose and fructose-containing sugars compared with dextrose (6–8). Differences in our findings may be due to the carbohydrate load used (300 kcals vs. 75 kcals in our study). Absorption of dextrose and fructose across small intestinal apical membranes occurs through sugar-specific primary transporters (24); however, high concentrations of either substrate can recruit GLUT2 to the apical surface, which has the capacity to transport both dextrose and fructose (25). This threshold for synergistic absorption could explain our lack of observation of elevated MR response to sucrose consumption. Similarly to others (6–8), we observed a lower CO response after dextrose consumption compared with sucrose and fructose. This attenuated response could be explained by the presence of fructose in the fructose and sucrose beverages and the differing metabolic fates of dextrose and fructose. While dextrose is primarily taken up for either storage or utilization for energy production by peripheral tissues (24), fructose is preferentially metabolized by the liver (26). In addition, fructokinase has a much greater affinity for fructose than glucokinase does for glucose (27), meaning it produces intermediates for further metabolism at a faster rate. Additionally, the first steps of fructolysis bypass the regulatory feedback mechanisms of glycolysis (28), including the inhibition of gluconeogenesis. Therefore, fructose metabolism can result in a futile cycle of glucose availability and oxidation in which glycolysis and gluconeogenesis occur concurrently (29).

### Bayesian Hierarchical Model

While both the traditional LMM and Bayesian Hierarchical Model analysis methods produced similar parameter estimates for the AUC, change from resting, and time to reach peak for CO, the Bayesian model resulted in reduced variance of these parameter estimates. Unlike the traditional LMM method, which first computes a single estimate for each individual observation and then fits a model to compare differences across conditions, the Bayesian model method first uses an assumed shape (cubic spline) to fit an average condition curve while estimating an individual participant’s curve. Then, the average condition curves are used to assess differences in parameters between groups, thus reducing the variability of the parameter estimate. In other words, when using the Bayesian model, smaller sample sizes can be used to elucidate physiological differences in metabolic responses. Given the well-documented inter-individual variability in assessments of postprandial MR and substrate oxidation (11, 30), this is an important feature of the model. Another distinguishing feature of this model is the capacity for borrowing strength, or the ability to pool the data across all observations and conditions to gain more knowledge about the parameters of interest (i.e., AUC, peak value, time at peak). Because individual observation curves are estimated simultaneously within the model, the model is robust to missing data; therefore, measurements that are varying lengths of time can still be fit in the model.

The Bayesian model has some limitations, however. Specifically, the model assumes data can be fit to a natural cubic spline; while this family of functions is reasonably flexible, it is possible that some participants’ data curves do not fit. In this regard, the LMM method is more versatile in that it does not assume a shape of the metabolic response curve. In addition, a direct computation of maximum values in metabolic response curves is not possible; a second model fit with the outcome measure as the absolute value of metabolic response would be required.

### Limitations

The current study has limitations. First, though our human study methods were designed to limit participant activity, we did not objectively measure movement inside the WRIC and cannot rule out the potential influence of movement on our measurements. Furthermore, while we removed data for the 4 minutes immediately following drink consumption, we cannot rule out persistence effects of movement on data beyond 4 minutes after drink consumption. Incorporating objective measures of activity will be an important component of future studies. Second, given our small WRIC volume, it is possible that body size could influence the volume derivative term used in our data post-processing calculations. While body volume likely does not affect data post-processing for WRIC studies in large-volume chambers, it is unclear whether this could be a meaningful source of measurement error in small-volume chambers. However, the within-subject crossover design of the present human study mitigates the potential influence of both participant movement and body volume on our results. Lastly, our sample was predominantly white, female, and had body mass indexes <25; as such, our results may not be generalizable to different populations.

### Conclusion and Future Applications

In summary, results from our technical and human studies highlight the reliability of our WRIC system in capturing steady state and dynamic metabolic measurements. Furthermore, our human study demonstrates the capacity to detect very small, dynamic changes elicited by small carbohydrate loads. These carbohydrate loads are more similar to those consumed in a serving of a sugar sweetened beverage than the 300 kcal loads used in previous research. Detection of these small post-ingestive changes is essential for understanding potential alterations of the gut-brain axis in people with obesity (31, 32). Finally, we proposed a new application for a statistical model, which can both estimate an individual curve for metabolic response for each participant within each condition and test for differences in key parameters of interest across conditions. This methodology, which is robust to missing data, reduces variance and can increase statistical power, potentially allowing for smaller sample sizes and therefore reduced cost for future studies. The validation and novel application of methods presented here provide important foundations for new research directions using WRICs to assess metabolic responses to small calorie loads.

## Acknowledgements

The authors would like to thank the research study participants who volunteered their time and efforts toward this project.

Validation data and de-identified human study data files, data dictionaries, and analysis code are made publicly available through the Virginia Tech Data Repository [doi to be placed here].

---

**Supplemental Figure 1.**
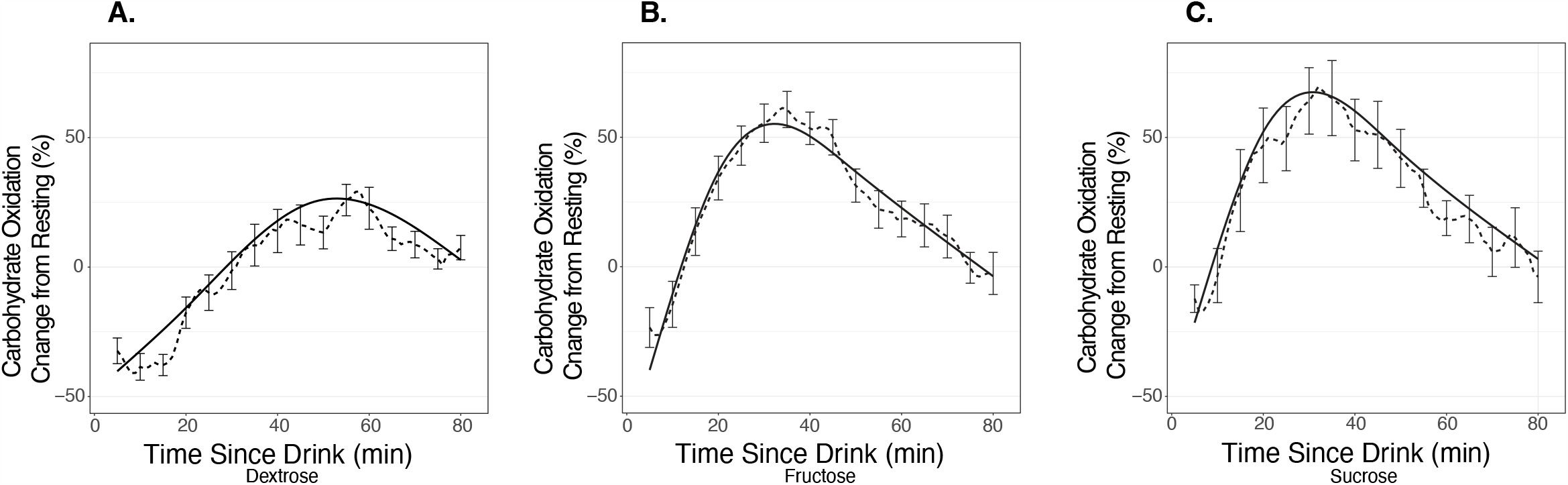
Observed postprandial change in carbohydrate oxidation (dashed lines) overlaid with Bayesian Hierarchical Model-estimated postprandial change in carbohydrate oxidation (solid lines) for (A) dextrose, (B) fructose, and (C) sucrose beverage conditions. Data are expressed as mean ± standard error of the mean for the observed changes.

**Supplemental Table 1:**
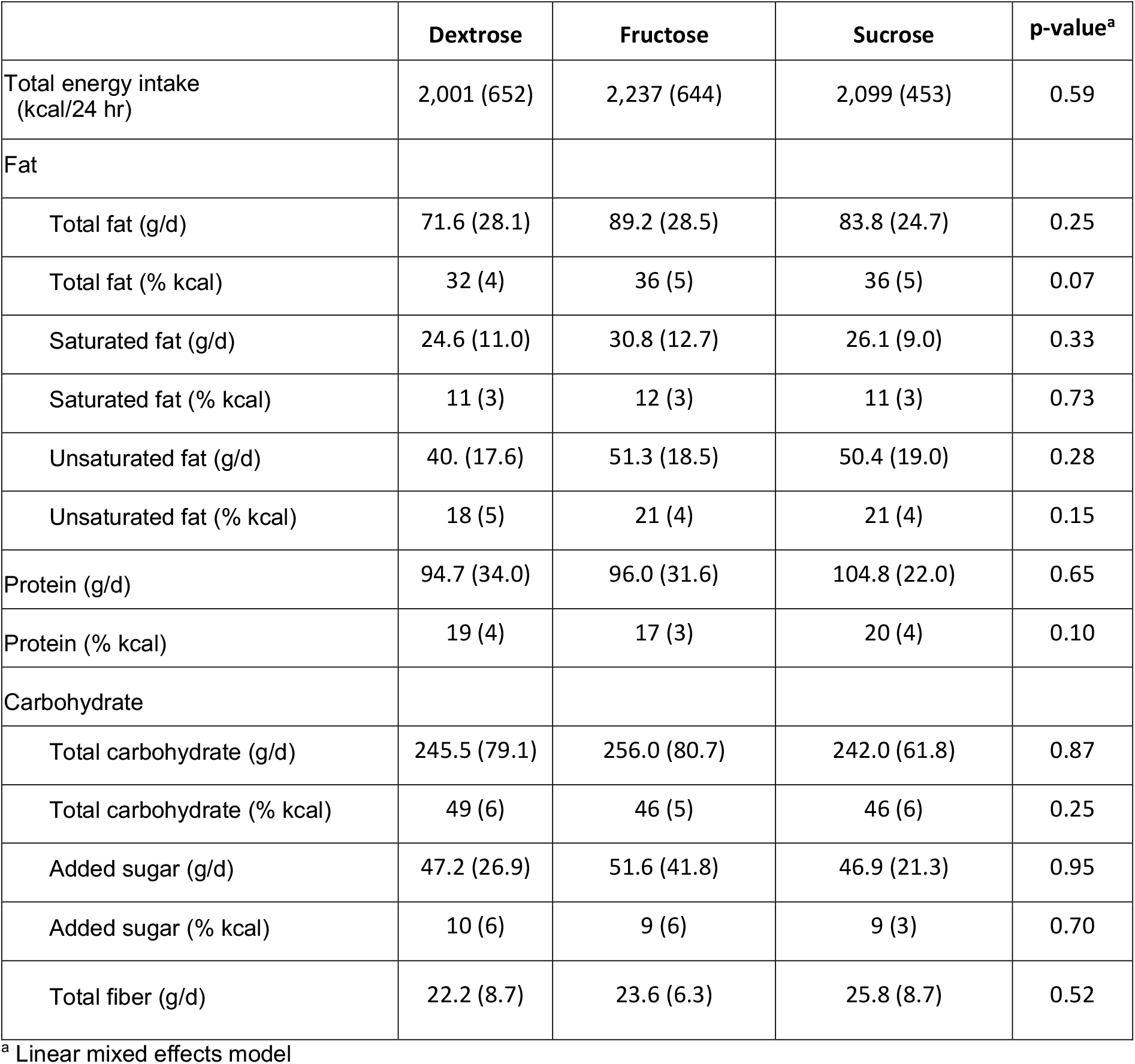
Dietary intake for 24 hours prior to each beverage condition indirect calorimetry session.

**Supplemental Table 2:**
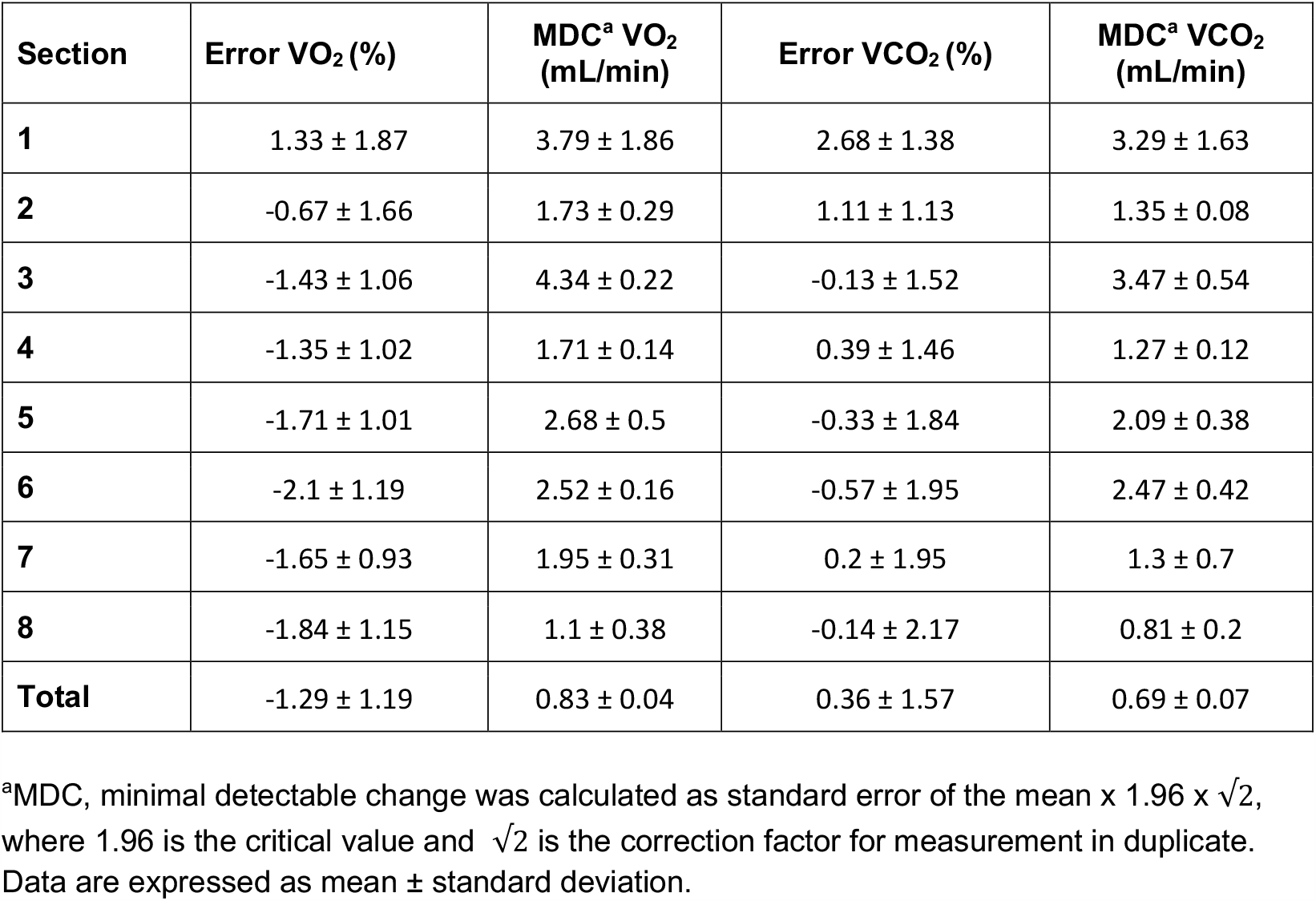
Reproducibility of O_2_ and CO_2_ recovery during infusion validation studies (n=4)

## References

1. Heinitz S, Hollstein T, Ando T, et al. Early adaptive thermogenesis is a determinant of weight loss after six weeks of caloric restriction in overweight subjects. Metabolism 2020;110:154303.

2. Garcia-Tabar I, Eclache JP, Aramendi JF, Gorostiaga EM. Gas analyzer’s drift leads to systematic error in maximal oxygen uptake and maximal respiratory exchange ratio determination. Frontiers in Physiology 2015;6.

3. Cooper JA, Watras AC, O’Brien MJ, et al. Assessing validity and reliability of resting metabolic rate in six gas analysis systems. J Am Diet Assoc 2009;109:128–132.

4. Chen KY, Smith S, Ravussin E, et al. Room Indirect Calorimetry Operating and Reporting Standards (RICORS 1.0): A Guide to Conducting and Reporting Human Whole-Room Calorimeter Studies. Obesity (Silver Spring) 2020;28:1613–1625.

5. Tappy L. Thermic effect of food and sympathetic nervous system activity in humans. Reprod Nutr Dev 1996;36:391–397.

6. Tappy L, Randin JP, Felber JP, et al. Comparison of thermogenic effect of fructose and glucose in normal humans. Am J Physiol 1986;250:E718–724.

7. Blaak EE, Saris WH. Postprandial thermogenesis and substrate utilization after ingestion of different dietary carbohydrates. Metabolism 1996;45:1235–1242.

8. Schwarz JM, Acheson KJ, Tappy L, et al. Thermogenesis and fructose metabolism in humans. Am J Physiol 1992;262:E591–598.

9. Schwarz JM, Schutz Y, Piolino V, Schneider H, Felber JP, Jéquier E. Thermogenesis in obese women: effect of fructose vs. glucose added to a meal. Am J Physiol 1992;262:E394–401.

10. Malik VS, Hu FB. The role of sugar-sweetened beverages in the global epidemics of obesity and chronic diseases. Nat Rev Endocrinol 2022;18:205–218.

11. Reed GW, Hill JO. Measuring the thermic effect of food. The American Journal of Clinical Nutrition 1996;63:164–169.

12. Schutz Y, Bessard T, Jéquier E. Diet-induced thermogenesis measured over a whole day in obese and nonobese women. Am J Clin Nutr 1984;40:542–552.

13. Conway JM, Ingwersen LA, Vinyard BT, Moshfegh AJ. Effectiveness of the US Department of Agriculture 5-step multiple-pass method in assessing food intake in obese and nonobese women. Am J Clin Nutr 2003;77:1171–1178.

14. Conway JM, Ingwersen LA, Moshfegh AJ. Accuracy of dietary recall using the USDA five-step multiple-pass method in men: An observational validation study. Journal of the American Dietetic Association 2004;104:595–603.

15. Weir JBDB. New methods for calculating metabolic rate with special reference to protein metabolism. J Physiol 1949;109:1–9.

16. Farinatti P, Castinheiras Neto AG, Amorim PRS. Oxygen Consumption and Substrate Utilization During and After Resistance Exercises Performed with Different Muscle Mass. Int J Exerc Sci 2016;9:77–88.

17. Péronnet F, Massicotte D. Table of nonprotein respiratory quotient: an update. Can J Sport Sci 1991;16:23–29.

18. de Valpine P, Turek D, Paciorek CJ, Anderson-Bergman C, Lang DT, Bodik R. Programming With Models: Writing Statistical Algorithms for General Model Structures With NIMBLE. Journal of Computational and Graphical Statistics 2017;26:403–413.

19. Dörner R, Hägele FA, Koop J, et al. Validation of energy expenditure and macronutrient oxidation measured by two new whole-room indirect calorimeters. Obesity 2022;30:1796–1805.

20. Thearle MS, Pannacciulli N, Bonfiglio S, Pacak K, Krakoff J. Extent and determinants of thermogenic responses to 24 hours of fasting, energy balance, and five different overfeeding diets in humans. J Clin Endocrinol Metab 2013;98:2791–2799.

21. Miles-Chan JL, Dulloo AG, Schutz Y. Fasting substrate oxidation at rest assessed by indirect calorimetry: is prior dietary macronutrient level and composition a confounder? Int J Obes (Lond) 2015;39:1114–1117.

22. Stinson EJ, Rodzevik T, Krakoff J, Piaggi P, Chang DC. Energy expenditure measurements are reproducible in different whole-room indirect calorimeters in humans. Obesity 2022;30:1766–1777.

23. Livesey G, Elia M. Estimation of energy expenditure, net carbohydrate utilization, and net fat oxidation and synthesis by indirect calorimetry: evaluation of errors with special reference to the detailed composition of fuels. Am J Clin Nutr 1988;47:608–628.

24. Stipanuk MH, Caudill MA. Biochemical, Physiological, and Molecular Aspects of Human Nutrition. Elsevier; 2018.

25. Gouyon F, Caillaud L, Carriere V, et al. Simple-sugar meals target GLUT2 at enterocyte apical membranes to improve sugar absorption: a study in GLUT2-null mice. J Physiol 2003;552:823–832.

26. ter Horst KW, Serlie MJ. Fructose Consumption, Lipogenesis, and Non-Alcoholic Fatty Liver Disease. Nutrients 2017;9:981.

27. Asipu A, Hayward BE, O’Reilly J, Bonthron DT. Properties of normal and mutant recombinant human ketohexokinases and implications for the pathogenesis of essential fructosuria. Diabetes 2003;52:2426–2432.

28. Merino B, Fernández-Díaz CM, Cózar-Castellano I, Perdomo G. Intestinal Fructose and Glucose Metabolism in Health and Disease. Nutrients 2019;12:94.

29. Chenoweth M, Dunn A. Fructose-6-phosphate substrate cycling and hormonal regulation of gluconeogenesis in vivo. Am J Physiol 1978;235:E295–303.

30. Weststrate J. Resting metabolic rate and diet-induced thermogenesis: a methodological reappraisal. The American Journal of Clinical Nutrition 1993;58:592–601.

31. Beutler LR, Corpuz TV, Ahn JS, et al. Obesity causes selective and long-lasting desensitization of AgRP neurons to dietary fat Elmquist JK, Dulac C (eds.). eLife 2020;9:e55909.

32. van Galen KA, Schrantee A, Ter Horst KW, et al. Brain responses to nutrients are severely impaired and not reversed by weight loss in humans with obesity: a randomized crossover study. Nat Metab 2023;5:1059–1072.

