## Supplemental Materials for "Validity and reliability of a new whole room indirect calorimeter to assess metabolic response to small-calorie loads"

**Supplemental Figure 1**

**Observed postprandial change in carbohydrate oxidation (dashed lines) overlaid with Bayesian Hierarchical Model-estimated postprandial change in carbohydrate oxidation (solid lines) for (A) dextrose, (B) fructose, and (C) sucrose beverage conditions. Data are expressed as mean ± standard error of the mean for the observed changes.**

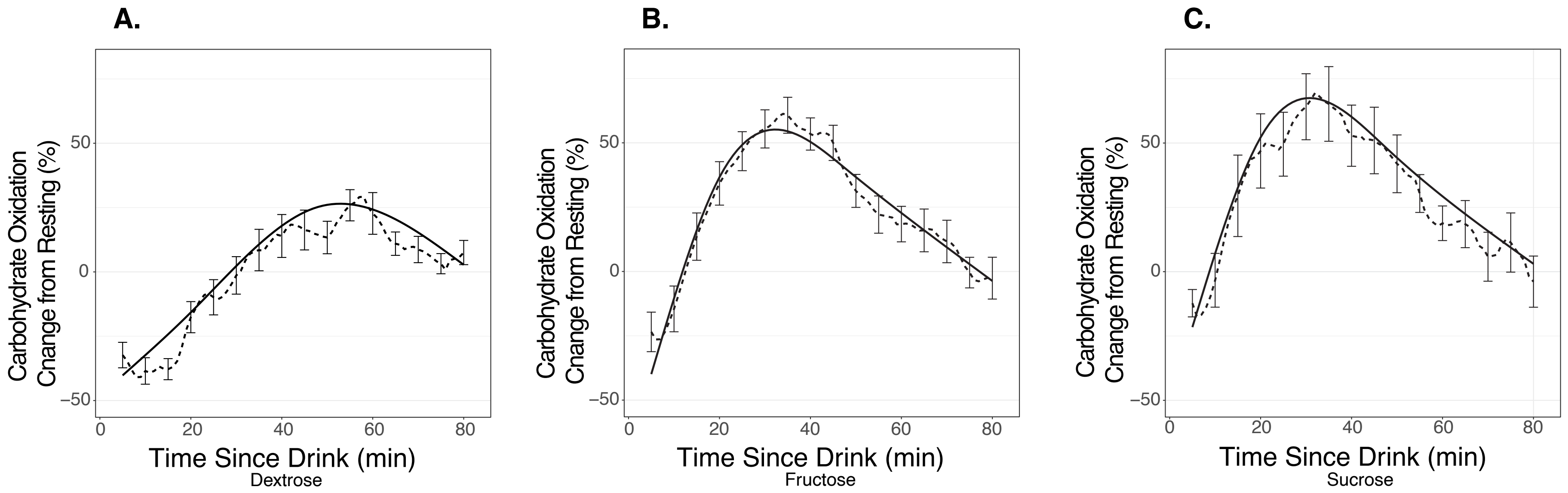

**Supplemental Table 1:** Dietary intake for 24 hours prior to each beverage condition indirect calorimetry session

| | **Dextrose** | **Fructose** | **Sucrose** | **p-value[a]** |
|---|---|---|---|---|
| Total energy intake (kcal/24 hr) | 2,001 (652) | 2,237 (644) | 2,099 (453) | 0.59 |
| Fat | | | | |
| Total fat (g/d) | 71.6 (28.1) | 89.2 (28.5) | 83.8 (24.7) | 0.25 |
| Total fat (% kcal) | 32 (4) | 36 (5) | 36 (5) | 0.07 |
| Saturated fat (g/d) | 24.6 (11.0) | 30.8 (12.7) | 26.1 (9.0) | 0.33 |
| Saturated fat (% kcal) | 11 (3) | 12 (3) | 11 (3) | 0.73 |
| Unsaturated fat (g/d) | 40. (17.6) | 51.3 (18.5) | 50.4 (19.0) | 0.28 |
| Unsaturated fat (% kcal) | 18 (5) | 21 (4) | 21 (4) | 0.15 |
| Protein (g/d) | 94.7 (34.0) | 96.0 (31.6) | 104.8 (22.0) | 0.65 |
| Protein (% kcal) | 19 (4) | 17 (3) | 20 (4) | 0.10 |
| Carbohydrate | | | | |
| Total carbohydrate (g/d) | 245.5 (79.1) | 256.0 (80.7) | 242.0 (61.8) | 0.87 |
| Total carbohydrate (% kcal) | 49 (6) | 46 (5) | 46 (6) | 0.25 |
| Added sugar (g/d) | 47.2 (26.9) | 51.6 (41.8) | 46.9 (21.3) | 0.95 |
| Added sugar (% kcal) | 10 (6) | 9 (6) | 9 (3) | 0.70 |
| Total fiber (g/d) | 22.2 (8.7) | 23.6 (6.3) | 25.8 (8.7) | 0.52 |

[a] Linear mixed effects model

**Supplemental Table 2:** Reproducibility of $O_2$ and $CO_2$ recovery during infusion validation studies (n=4)

| **Section** | **Error $VO_2$ (%)** | **MDC[a] $VO_2$ (mL/min)** | **Error $VCO_2$ (%)** | **MDC[a] $VCO_2$ (mL/min)** |
|---|---|---|---|---|
| **1** | 1.33 ± 1.87 | 3.79 ± 1.86 | 2.68 ± 1.38 | 3.29 ± 1.63 |
| **2** | -0.67 ± 1.66 | 1.73 ± 0.29 | 1.11 ± 1.13 | 1.35 ± 0.08 |
| **3** | -1.43 ± 1.06 | 4.34 ± 0.22 | -0.13 ± 1.52 | 3.47 ± 0.54 |
| **4** | -1.35 ± 1.02 | 1.71 ± 0.14 | 0.39 ± 1.46 | 1.27 ± 0.12 |
| **5** | -1.71 ± 1.01 | 2.68 ± 0.5 | -0.33 ± 1.84 | 2.09 ± 0.38 |
| **6** | -2.1 ± 1.19 | 2.52 ± 0.16 | -0.57 ± 1.95 | 2.47 ± 0.42 |
| **7** | -1.65 ± 0.93 | 1.95 ± 0.31 | 0.2 ± 1.95 | 1.3 ± 0.7 |
| **8** | -1.84 ± 1.15 | 1.1 ± 0.38 | -0.14 ± 2.17 | 0.81 ± 0.2 |
| **Total** | -1.29 ± 1.19 | 0.83 ± 0.04 | 0.36 ± 1.57 | 0.69 ± 0.07 |

[a]MDC, minimal detectable change was calculated as standard error of the mean x 1.96 x $\sqrt{2}$, where 1.96 is the critical value and $\sqrt{2}$ is the correction factor for measurement in duplicate.
Data are expressed as mean ± standard deviation.
